# Probing Nanoscale Electrochemical Phenomena with Nitrogen-Vacancy Centers in Diamond

**DOI:** 10.1101/2024.11.29.626096

**Authors:** Samuel Fulton, Jack Stropko, Robert Vitale, Alexander O. Sushkov

## Abstract

Nitrogen-vacancy centers in nanodiamonds exhibit photo-stability and biocompatibility that make them promising candidates for versatile biological sensors. In the present work, we study the dependence of NV-nanodiamond fluorescence on the pH of the surrounding ionic aqueous solution. Band-bending effects and modified ion migration rates may be the potential mechanisms underlying the observed pH sensitivity. Our work offers insight into diamond electrochemistry and paves the way toward nanoscale pH imaging. Additionally, the methodologies developed in this work introduce a viable approach for analyzing local electrochemical environments, with potential applications in intra-cellular pH sensing, the design of electrolytic cells, and the development of alternative fuel technologies.

## INTRODUCTION

Nitrogen-vacancy (NV) centers in diamond, consisting of a nitrogen impurity and neighboring lattice vacancy, are one of many fluorescent centers used in bio-sensing [1]. The spin-dependent fluorescence of the NV center is sensitive to external factors such as magnetic field, temperature, lattice strain, and electric field, making NV centers excellent environmental sensors [2, 3]. Because of NV centers’ long spin coherence times and lack of photo-bleaching, they have been extensively employed in quantum computation, quantum metrology, and spintronics [4–8].

In this study, we diverge from conventional spin-based measurements and instead probe voltage-induced fluorescent modulation. Specifically, we investigate the fluorescent modulation that arises from the switching of NV centers in nanodiamonds (NDs) between the neutral state (NV^0^) and the negatively charged state (NV^*−*^) [9, 10]. The fluorescence spectra of both charge states are shown in Fig. 1. Under 532 (nm) excitation, the NV^*−*^ fluoresces more intensely than the NV^0^ for wavelengths longer than 630 (nm), as shown in Fig. 1. We observe voltage-induced charge state switching by measuring changes in fluorescence, where an increase in fluorescence corresponds to NVs moving from the NV^0^ state to the NV^*−*^ state.

**FIG. 1.**
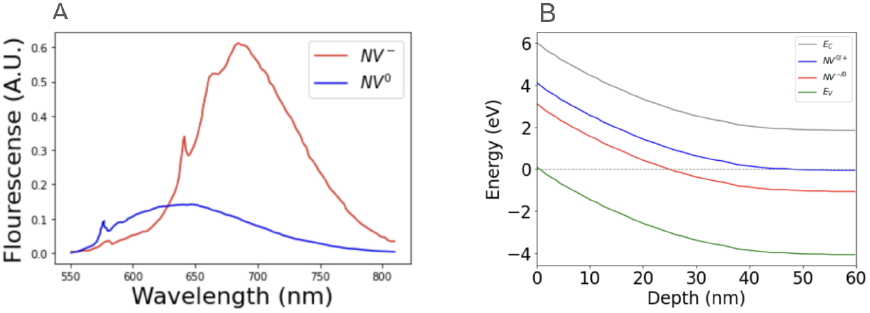
(A) The normalized fluorescence spectra of NV^0^ and NV^*−*^, retrieved from Ref. [11]. To preferentially detect fluorescence from NV^*−*^, we use a 665 nm long-pass filter. (B) The electronic band structure of NV centers in hydrogen-terminated nanodiamonds, retrieved from Ref. [12]. By applying a voltage to the electrochemical cell we shift the Fermi level causing switching between the NV^0^ and NV^*−*^ charge states.

We observe that the voltage-induced charge state switching depends on the pH of the electrochemical cell, when the pH is varied between 7 and 12. A possible explanation for this change in behavior is a change in the electron band structure of NDs. Changes in ion migration rates inside the solution offer another possible explanation for the change in voltage response at high pH. Regardless of the mechanisms, this work offers interesting applications in nanoscale pH sensing.

## MATERIALS AND METHODS

We use a standard NV optical setup that includes a scanning confocal microscope equipped with an inverted oil-immersion objective, Fig. 2. We focus a 532 nm continuous-wave laser on the back aperture of the objective. Under 532 nm excitation, the NV^*−*^ fluoresces more intensely than the NV^0^ for wavelengths longer than 630 nm, as shown in Fig. 1. We use a 665 nm cutoff long-pass filter to preferentially detect fluorescence from NV^*−*^. We collect the fluorescence signal from NV centers with the same objective and focus the beam through a series of lenses onto an avalanche photodiode.

**FIG. 2.**
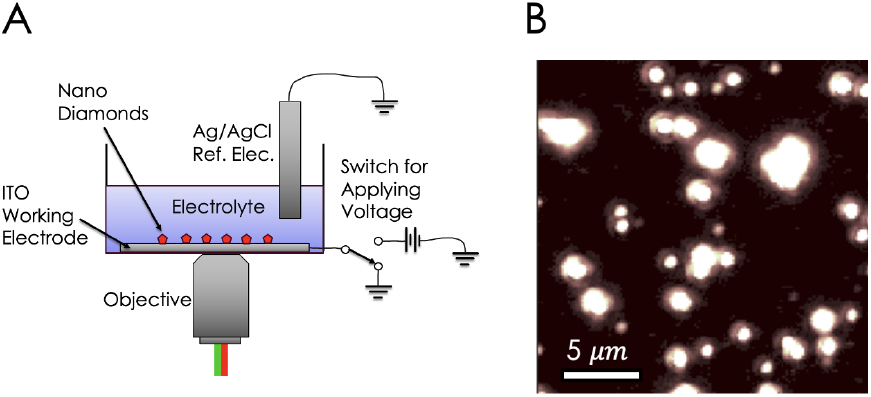
(A) Nanodiamonds are at the bottom of the petri dish, on an ITO conducting electrode. The pH of the electrolyte solution is altered and fluorescence voltage response is measured. The switch is normally open, grounding the ITO working electrode. A voltage pulse is applied by closing the switch. The Ag/AgCl reference electrode is always grounded. (B) The fluorescence intensity, mapped using the confocal microscope. The bright spots are either nanodiamonds or clusters of nanodiamonds.

The electrochemical cell consists of a petri dish with a hole in its base, sealed with a coverslip coated with indium-tin-oxide (ITO). The ITO is applied to the coverslip via sputtering. Hydrogenated nanodiamonds (NDs) are deposited on the coverslip by pipetting an aqueous solution and allowing it to dry. After drying, the NDs are securely attached to the ITO through electrostatic interactions. The mean diameter of the NDs is 100 nm.

A copper wire leading to a signal generator is epoxied to the coverslip such that a voltage can be applied to the ITO and NDs. The petri dish is filled with an aqueous electrolyte solution consisting of 10 mM sodium chloride (NaCl). The voltages are applied across the ITO working electrode and a silver/silver chloride (Ag/AgCl) reference electrode, which is grounded. The reference electrode material was chosen because of its stability for a wide range of pH values. Applying a voltage across the circuit results in a voltage drop across the Helmholtz double layer at the ITO-electrolyte interface. The applied voltage causes shifts in the Fermi level and results in NV charge state switching. The charge state switching is observed by changes in fluorescence. Charge state switching is well documented in Ref. [13]. To study electrochemical processes, we apply voltage pulses to the electrochemical cell at differing solution pH values. Specifically, we conduct measurements at pH 7, achieved using DI water and an electrolyte solution, and at pH 12, obtained by adding a concentrated pH 13 KOH solution. Interestingly, the voltage-dependent fluorescent response changes with pH.

The applied voltage is defined as

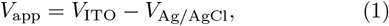

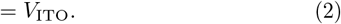

## MEASUREMENT

We apply a voltage pulse for 1.5 seconds and record the resulting change in fluorescence. For each time series trace, we collect 40 averages, over the experimental timescale of roughly 3 minutes. At the pH values of 7 and 12, we characterize fluorescent response by incrementally adjusting the pulse voltage amplitude from -3 V to 3 V, as shown in Fig. 3. To better quantify the size of the voltage response, we take the integral of the fluorescence time series data and display the results in Fig. 3.

**FIG. 3.**
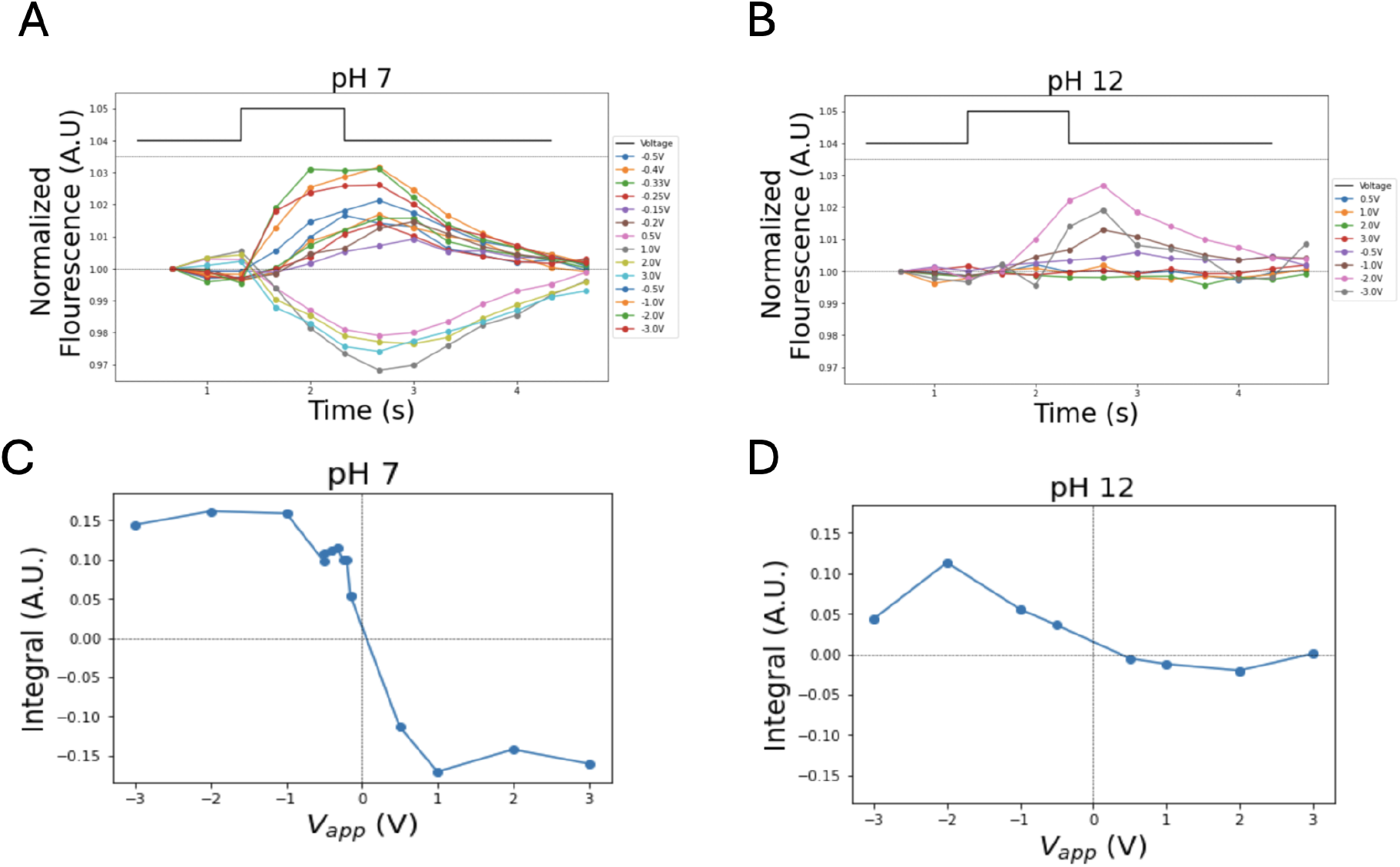
(A) At the solution pH of 7, a voltage *V*_app_ is applied for 1.5 seconds, indicated by the square pulse. The fluorescence time series data are shown for voltage pulses ranging from *−*3 V to 3 V. (B) The process from panel (A) is repeated at the pH of 12. (C and D) The time-integral of the voltage response was calculated at pH of 7 and 12 and plotted as a function of applied voltage.

**FIG. 4.**
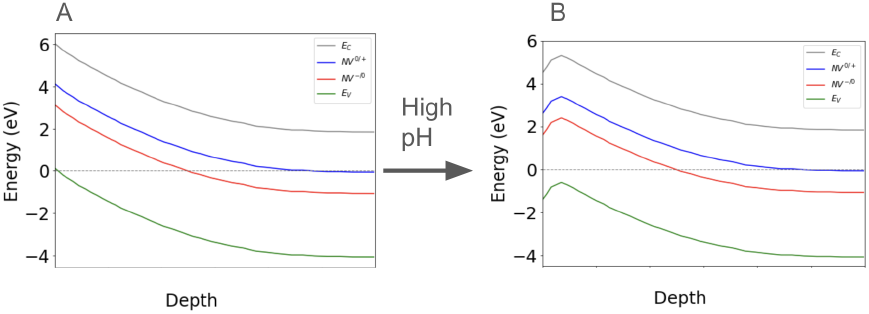
A pictorial representation of pH-induced downward band bending, following Ref. [16]. (A) The hydrogenated diamond band structure at pH of 7. (B) Band structure at pH of This phenomenon was used to explain the disappearance of 2DHG (p-type surface doping) in Ref. [16]

At the pH of 7, applying a negative bias increases fluorescence while applying a positive bias decreases fluorescence, consistent with the results in Ref. [13]. However, after increasing the pH to 12 by adding a solution of KOH (pH 13), the response to positive *V*_app_ is significantly reduced, while a degree of sensitivity to negative *V*_app_ remains. The pH-dependent asymmetry presents potential applications in nanoscale pH imaging and insight into nanoscale electrochemical processes.

The time scale of the fluorescence response is set by the charging of the ITO/electrolyte interface. This is similar to the charging of a capacitor, but rather than an RC time constant, the time scale is set by the migration rate of charged ions in the solution [14]. We note that since measurements shown in Fig. 3 include data taken with voltage pulses of 1.5 seconds, slower diffusion dynamics may be obscured. However, this does not negate the potential utility of nanoscale pH measurements performed here [15].

## MECHANISM

We propose two possible mechanisms for the change in behavior at high pH. The first mechanism is based on adsorbed-ion-induced band bending, which provides a means of screening the effect of applied voltages. When a positive voltage bias is applied, negative ions (Cl^*−*^ and OH^*−*^) migrate towards the working electrode. The localized increase in OH^*−*^ concentration at the diamond surface enhances downward band bending in the nanodiamonds, consistent with the mechanisms outlined in Ref. [16]. The increase in downward band bending counters the decrease in Fermi level, screening any decrease in fluorescence.

Another possible explanation for the reduction of voltage response comes from a shift in ion migration rates. Longer time scales are likely set by ion migration dynamics [14]. Ongoing measurements are investigating the fluorescence response on much longer time scales of tens of seconds.

## APPLICATIONS

Regardless of uncertainties in underlying mechanisms, the nanoscale pH measurements offer a wide range of applications. Intracellular pH is a key parameter that influences many biochemical and metabolic pathways and can be an indirect marker of neural activity and cell health [17]. Unlike previous NV pH measurements, which require either small nanodiamonds (10-20 nm diameter) or long averaging times (multiple hours), our measurements work for large nanodiamonds (100 nm diameter) and require less than 5 minutes of averaging [1, 18]. Furthermore, our technique is well suited for studying nanoscale dynamics in solution and measurements of solution pH fluctuations.

Our experimental setup is quite similar to a water electrolysis experiment. In water electrolysis, the application of a voltage across the electrodes produces oxygen (O_2_) and hydrogen (H_2_) gas. Water electrolysis is of paramount importance for the development of alkaline electrolyzers and alkaline fuel cell (AFC) systems [19]. It has been known for decades that electrolyte pH affects the current across the electrodes by orders of magnitude. However, the origins of this effect are still under debate [20]. Our measurement may have the potential to offer insight into the mechanism of the effect of pH on electrolytic activity, which is a step toward alternative fuel sources.

## OUTLOOK

Our work studies how NV charge state in NDs responds to an applied voltage in an electrochemical cell. We demonstrate that charge state switching depends on the pH of the solution. While the underlying mechanism is still uncertain, the results of this study offer potential applications in biology and chemistry. This measurement paves the way toward nanoscale and intracellular pH measurements. Furthermore, the results of our study offer a new method for studying electrocatalysts, which may be key in the development of renewable fuel sources.

We are conducting a series of experiments to uncover the fundamental mechanisms. Our current focus is on the dynamical timescales, which are governed by diffusion rates. In parallel, we are measuring the voltage response under acidic pH conditions, with plans to expand this investigation by varying NaCl concentrations across different pH levels. Additionally, we aim to study how the NV center spin *T*_1_ and *T*_2_ relaxation times depend on the surrounding solution pH. Of interest is the behavior at pH levels above 11, where hydrogen-terminated nanodiamonds are predicted to lose surface conductivity [21]. This loss is expected to affect relaxation rates.

